# Life Cycle and Morphogenetic Differentiation in Heteromorphic Cell Types of a Cosmopolitan Marine Microalga

**DOI:** 10.1101/2024.07.03.601694

**Authors:** Laurie Bousquet, Shai Fainsod, Johan Decelle, Omer Murik, Fabien Chevalier, Benoit Gallet, Rachel Templin, Yannick Schwab, Yoav Avrahami, Gil Koplovitz, Chuan Ku, Miguel J. Frada

## Abstract

- *Gephyrocapsa huxleyi* is a prevalent, bloom-forming phytoplankton species in the oceans. It exhibits a complex haplo-diplontic life cycle, featuring a diploid-calcified phase, a haploid phase, and a third ’decoupled’ phase produced during viral infection. Decoupled cells display a haploid-like phenotype, but are diploid.
- Here, we investigated the fate of decoupled cells during culture observations and we compared the transcriptome profiles and the cellular ultrastructure of the three cell types.
- We found that decoupled cells can revert to the calcified form in the absence of viral pressure, revealing the transient nature of this cell type. Ultrastructural analyses showed distinct nuclear organisation with variations in chromatin volume. Transcriptomic analyses revealed gene expression patterns specific to each life phase. These included multiple regulatory functions in chromatin remodelling, broader epigenetic mechanisms and life cycling, which likely contributed to cell differentiation. Finally, the exploration of available host-virus transcriptomes supports life cycle transition during viral infection.
- This study provides cellular and molecular foundations for nuclear remodelling and cell differentiation in coccolithophores and the identification of gene markers for studying coccolithophore life cycles in natural populations.

## Introduction

Eukaryotes exhibit complex life cycles comprising multiple phases interlinked by sexual mechanisms (syngamy and meiosis). They involve gene recombination, variations in ploidy levels (haploid,1n and diploid, 2n), as well as asexual mechanisms of cell differentiation. These transitions are crucial for the developmental and reproductive cycles of organisms, as well as for the adaptation to environmental changes through the differentiation of physiologically fit or resistant phenotypes (Mable & Otto, 1998; Von Dassow & Montresor, 2011).

Coccolithophores are a prevalent group of marine unicellular algae that produce distinct CaCO_3_ plates, coccoliths, coating the cell surface (Monteiro *et al*., 2016). Due to photosynthesis and calcification, these microorganisms play a pivotal role in the global carbon cycle (Balch, 2018). Coccolithophores have complex sexual haplodiplontic life cycles, composed of 1n- and calcified 2n-cell phases, each capable of independent cell divisions and expressing dissimilar morphology (heteromorphic). Such life cycles are proposed to widen niche-space and facilitate the adaptation to environmental variability (Houdan *et al*., 2004; Frada *et al*., 2019; de Vries *et al*., 2021).

*Gephyrocapsa huxleyi* (formerly *Emiliania huxleyi*, (Bendif *et al*., 2019), is the most prevalent coccolithophore in the ocean and forms extensive seasonal blooms in high latitude regions, impacting marine ecology and biogeochemistry (Balch, 2018).The haplodiplontic, heteromorphic life cycle of *G. huxleyi* features diploid calcified cells (2n-calcified) that are non-motile and haploid non-calcified, biflagellated cells, coated by simpler organic scales (1n-flagellated) (Klaveness, 1972; Green *et al*., 1996).

Although 1n cells have been detected in 2n cultures, controlled manipulation of sexual transitions (syngamy or meiosis) has not yet been achieved. Transcriptome analyses have identified vast differences between diploid and haploid cells, particularly in transcripts linked to calcification, membrane trafficking and motility genes expressed in 2n-calcified cells and 1n-flagellated cells, respectively (von Dassow *et al*., 2009; Rokitta *et al*., 2011; Rokitta & Rost, 2012). However, despite extensive research on the ecological and biogeochemical roles of the 2n-calcified cells, the ecology of 1n-flagellated cells remains poorly understood (Frada *et al*., 2012).

A major influence on *G. huxleyi* blooms are specific lytic viruses (*Emiliania huxleyi*- specific viruses, EhV) that infect diploid cells and drive bloom decay (e.g. Bratbak *et al*., 1993; Vardi *et al*., 2012). Strikingly, culture experiments have demonstrated that 1n cells are resistant to EhV (Frada *et al*., 2008). In addition, during infection of 2n- calcified cells, cells displaying a haploid-like phenotype (biflagellated, organic scale-bearing) and resistant to EhV are produced during viral infections (Frada et al. 2017). These cells are however diploid, thus referred to as the ’decoupled ’ cell type (2n- flagellated) (Frada *et al*., 2017), whose production resembles apospory reported in plants and algae, where diploid gametophytes (typically haploid) develop without meiosis (Coelho *et al*., 2007). Such differentiation of 2n-flagellated cells may confer an escape strategy from EhV, potentially setting the stage for post-bloom recovery (Frada *et al*., 2017).

Life cycle transitions typically involve layers of mechanisms, notably epigenetic processes. These include chemical modifications of chromatin via the addition or removal of chemical moieties on histone tails (histone post-translational modifications, hPTMs) driving chromatin ultrastructural remodelling and gene expression regulation (Lawrence *et al*., 2016). Some hPTMs and associated proteins are implicated in life cycle regulation. Notably, the Polycomb Repressive Complex 2 (PRC2) mediates the silencing of key three-amino acids loop extension (TALE) homeodomain transcription factors (KNOX- and BELL-like) that in turn, regulate haploid-to-diploid transitions in land plants (Sakakibara *et al*., 2013; Horst *et al*., 2016; Dierschke *et al*., 2021) and in multiple algae (e.g., Lee *et al*., 2008; Thangavel & Nayar, 2018; Arun *et al*., 2019; Mikami *et al*., 2019; Hirooka *et al*., 2022).

Specificities in these mechanisms are associated to different species. However, alterations of these life cycle regulators can result in the decoupling of ploidy and life phase generation (Schmidt, 2020), underlying either apospory (defined above) or apogamy where instead haploid sporophytes (typically diploid) develop without gamete fusion (e.g., Zhao *et al*., 2001; Coelho *et al*., 2011; Sakakibara *et al*., 2013)

Here, we surveyed the fate of 2n-flagellated (decoupled) cells in culture of the cosmopolitan microalga *G. huxleyi*. Moreover, we examined their relation to 2n- calcified and 1n-flagellated forms, from morphological and gene expression perspectives using 3D-microscopy (FIB-SEM, Focused ion beam scanning electron microscopy) and transcriptome analyses. Finally, we also probed published *G. huxleyi*-EhV transcriptomes to examine 2n-flagellated cell differentiation during viral infection. Our results provide novel insights on the regulatory mechanisms driving life cycle differentiation and response to viruses in *G. huxleyi*.

## Material and Methods

### Algal strains, maintenance and viral infections

We used *Gephyrocapsa huxleyi* strains RCC1216 (2n-calcified, Houdan *et al*., 2005), LC5-11A (1n-flagellated), and LC5-12A (2n-flagellated) in this study. LC5-11A was isolated from RCC1216 via single-cell pipetting following a spontaneous life cycle change in culture, 6 months prior the study. LC5-12A and other 2n-flagellated strains were isolated from RCC1216 during an EhV 201 infection (Schroeder *et al*., 2002) 4 to 6 months prior the study, as described in Frada *et al*., 2017. Cultures, maintained in K/2 (-Tris, -Si) medium (Keller *et al*., 1987), were not axenic. Genome sizes were validated by flow cytometry of isolated nuclei (von Dassow *et al*., 2009). Ten additional decoupled strains were isolated from 3 independent RCC1216 infection experiments with EhV 201. Over 2 years, routine inspection of decoupled cultures was performed using optical microscopy. Sensitivity to EhV201 infection was assessed during the exponential growth phase with a multiplicity of infection of 0.2. Cell and viral concentrations were determined by flow cytometry (Attune NxT Life Technology).

### Algal growth for transcriptome analyses

The 3 strains were grown in triplicate in 4L borosilicate vessels, under gentle air-bubbling at 18°C, illuminated with 80 µmol photons m⁻²s⁻¹ from warm-white LED panels on a 14:10h light/dark cycle. Cultures were initiated at 5-8 x 10⁴ cells mL⁻¹ and monitored for 48h by flow cytometry. The maximum photochemical quantum yields of PSII (Fv/Fm), were measured by fluorescence induction and relaxation system (Satlantic Inc., Halifax, NS, Canada; Gorbunov and Falkowski, 2005) (Table S1). Four hours after dawn, 200 mL of each culture was harvested and pelleted by centrifugation (8000 g, 10 min, 4°C). Cell pellets were flash-frozen in liquid N₂ and stored at -80°C until RNA extraction.

### RNA isolation, libraries preparation and sequencing

RNA was extracted from cell pellets using the RNeasy Plant Kit (Qiagen) and treated with DNase (Turbo DNA-free kit, Ambion). RNA concentration and integrity were measured by Nanodrop (Thermo Fisher Scientific) and Tapestation (Agilent Genomics). cDNA libraries were prepared according to the Illumina RNA sample preparation kit. mRNA was isolated using poly(T) oligo-attached magnetic beads, fragmented into 200-500 bp pieces and RNA fragments were reverse transcribed with SuperScript II reverse transcriptase (Life Technology). Adapters ligation, amplification and purifications preceded the sequencing of the cDNA libraries on an Illumina NextSeq platform into 75-bp single-end reads.

### Transcriptome curation, assembly and annotation

FastQC (www.bioinformatics.babraham.ac.uk) and Trimmomatic (Bolger *et al*., 2014) were used for quality assessment and adapter trimming. Reads from the 9 *G. huxleyi* libraries (triplicates for RCC1216, LC5-10A, LC5-12A) were aligned to the *G. huxleyi* RCC1217 genome (Kao *et al*., 2024), a haploid strain isolated from RCC1216.

Alignment was performed with STAR v2.7.10a (Dobin *et al*., 2013). Transcripts were reconstructed and merged into a synthetic life phase transcriptome using StringTie v2.2.1 (Pertea *et al*., 2015), Cuffmerge v2.2.1 (Trapnell *et al*., 2012), and gffread v0.12.7 (Pertea & Pertea, 2020), yielding 60930 transcripts. Transcriptome completeness was evaluated with BUSCO v5.2.2 (Manni *et al*., 2021) against the eukaryota_odb10 database. Redundancy was reduced with CD-HIT-EST v4.8.1 (Fu *et al*., 2012), resulting in 56092 non-redundant transcripts. Coding sequences were identified with TransDecoder v5.5.0 . Functional annotation combined InterProScan v5.52-86.0, Diamond blastP v2.0.14.152 against the nr database and EggNOG mapper (Huerta-Cepas *et al*., 2019; Blum *et al*., 2021; Buchfink *et al*., 2021). Results were consolidated in OmicsBox, and Gene Ontology annotations were retrieved using Blast2GO (Götz *et al*., 2008). Parameters are detailed in Methods S1, life phase transcriptome information in Methods S2.

Histones, chromatin associated transcripts (including transcription factors) were examined using two complementary strategies: (1) identification of close homologs through Blast similarity searches against local databases and (2) detection of conserved protein domains with HMMER v3.3.2. Only shared hits were retained for subsequent analyses. Databases were assembled from Thiriet-Rupert *et al*., 2016; Grau-Bové *et al*., 2022 and https://planttfdb.gao-lab.org. Details are provided in Methods S3.

### Transcriptome Quantification, Differential Expression and Clustering

RSEM v1.3.3 (Li & Dewey, 2011) with the parameter --bowtie2 was used for gene-level abundance estimation. Features with fewer than 3 raw counts in at least 3 samples were filtered out. Transcripts per Million (TPM) normalization was applied, and features with a mean expression > 0.01 TPM were considered expressed.

Differentially expressed transcripts were identified using DESeq2 (Love *et al*., 2014), with those showing a log2fold change > |2| and FDR < 0.05 clustered using K-means. The optimal number of clusters was determined through complementary statistical methods and visual inspection for homogeneity and separation. The same approach was applied to chromatin associated transcripts, with a selection of log2fold change > |0.5| and subsequent clustering.

Enrichment analysis was performed in R (v4.2.2; R Core Team 2021) using the hypergeometric (HG) test. Null p-values were adjusted by adding an epsilon value (smallest non-zero p-value divided by 1000) and applying the Benjamini-Hochberg (BH) correction (Benjamini & Hochberg, 1995). Data analysis and visualization were conducted in R, Unix and Python. Software parameter settings are detailed in Methods S1. Validation of differential expression between cell types was performed by qRT-PCR using various life phase-specific genes (Methods S4, Table S2).

### Analyses of available *G. huxleyi* transcriptomes during viral infection

Data from studies on the transcriptomic dynamics of *G. huxleyi* during viral infection (Feldmesser *et al*., 2014; Rosenwasser *et al*., 2014) were used to examine the expression patterns of specific life phase transcripts. *G. huxleyi* CCMP2090 was infected with lytic viral strain EhV 201 at a multiplicity of infection of ∼1:1. Samples were collected at 1h and 24h post-infection, with non-infected cultures as controls, as indicated in the original studies. In the present study, life phase transcripts were blasted against CCMP2090 transcripts to identify homologs, retaining those with 100% identities over 80% alignment length. We focused on "flagellated-specific" transcripts (expressed exclusively in flagellated forms - haploid and decoupled cells). Abundance estimations from CCMP2090 were used to track the expression of these transcripts over the infection timeline (see Table S3 for functional annotation and clustering information of CCMP2090 homologs to flagellated transcripts).

### Focused Ion Beam-Scanning Electron Microscopy (FIB-SEM): sample preparation and analysis

*G. huxleyi* life stages were maintained under similar conditions as described above and harvested during exponential growth. Cells were centrifuged at 3000 g for 2 min, cryo-fixed using high-pressure freezing (HPM100, Leica), and freeze-substituted (EM ASF2, Leica) as in Decelle *et al*., 2019; Uwizeye *et al*., 2021 and Gallet *et al*., 2024. Samples were embedded in 100% resin, mounted on SEM stubs, gold sputter-coated (Quorum Q150RS; 180 s at 30 mA), and imaged with a FIB-SEM (Crossbeam 540, Carl Zeiss). Atlas3D software (Fibics Inc. and Carl Zeiss) was used for sample preparation and 3D acquisitions, with SEM images recorded at an 8 nm pixel size. Segmentation of organelles (plastids, mitochondria, nucleus) and cellular compartments was performed using 3D Slicer software (Oyama *et al*., 2014) using a manually-curated, semi-automatic pixel clustering mode (Decelle *et al*., 2021; Uwizeye *et al*., 2021). Colors were assigned to segmented regions, and image intensity thresholds were adjusted. Morphometric analyses were conducted with the "segmentStatistics" module in 3D Slicer, converting values to µm³ or µm² based on an 8 nm voxel size. Five diploid-calcified cells, five haploid cells, six 2n-flagellated cells (non-calcified), and four reverted calcifying cells were analysed. Further methodological details are provided in Methods S5.

## Results

The life cycle of *G. huxleyi* includes three cell types representing distinct phases: 2n- calcified (RCC1216), 1n-flagellated (LC5-10A) and 2n-flagellated (LC4-12A) cell type (Fig. 1a). Nuclear DNA content analyses by flow cytometry confirmed the ploidy of the three cell-types (Fig. S1a). The fate of decoupled cells and the morphogenetic differentiation between the three cell types were examined in our study.

**Figure 1.**
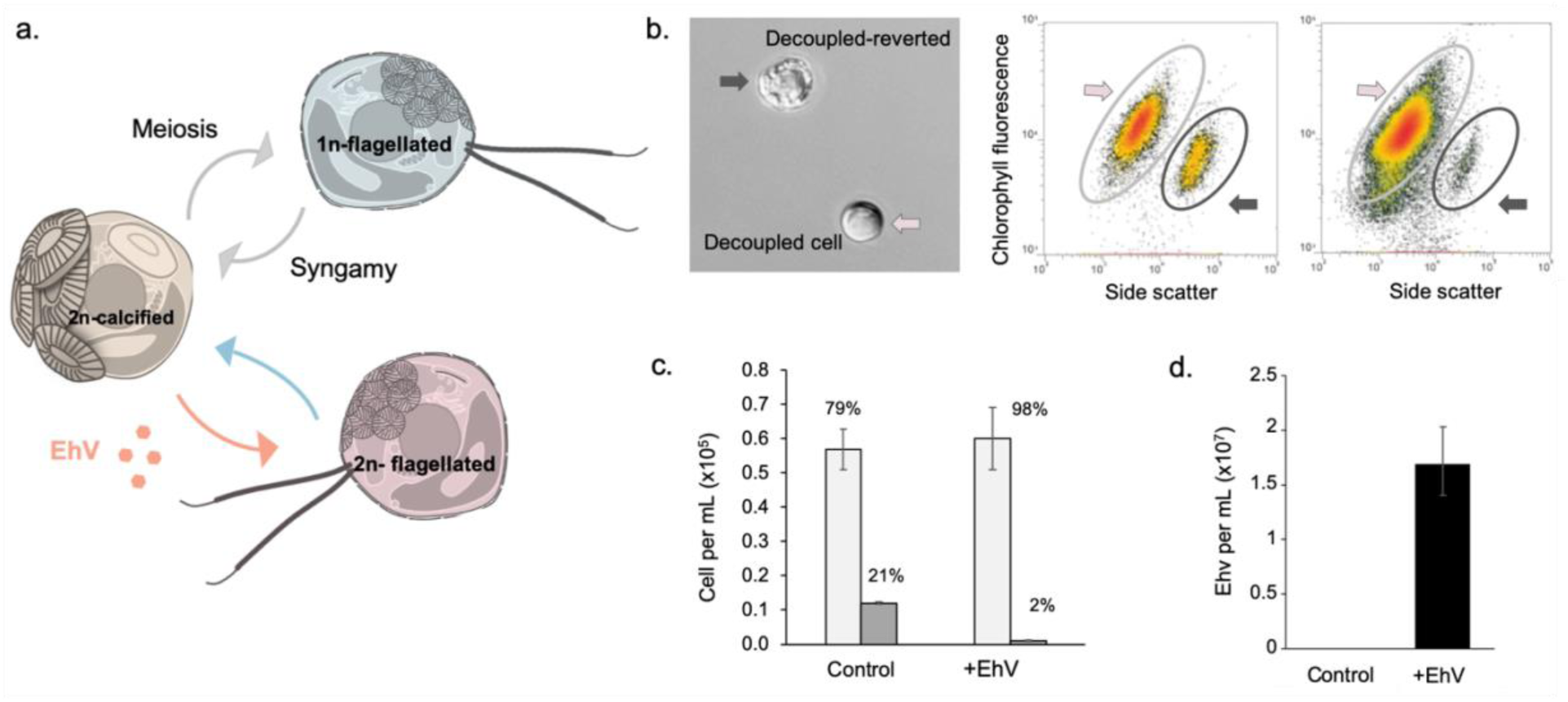
*Gephyrocapsa huxleyi* life cycle, reversion and response to viral infection. (a) Schematic illustration of the life cycle of *G. huxleyi* and role of virus on life cycle transitions. Sexual reproduction (meiosis and syngamy) drives the alternation between haploid, uncalcified, biflagellated cells (1n-flagellated, grey) and diploid calcified cells (2n-calcified, orange). During infection by specific *Emiliania huxleyi* viruses (EhV), 2n-calcified cells can produce 2n-flagellated cells (red) that are phenotypically similar to 1n-flagellated cells. This phenotype is reversible in absence of viral pressure (see results). 1n and 2n: haploid and diploid respectively (b) Microscopy and flow cytometry depicting the emergence of calcified cells in 2n-decoupled cultures in the absence of EhV. Left panel: light microscopy images of a 2n-flagellated (light red arrow) and reverted (dark grey arrow) cells, co-isolated in a clonal culture of 2n-flagellated cells in absence of viral pressure. Central and right panels: Gating of uncalcified, bi-flagellated (light grey ellipses) and calcified populations (reverted cells, dark grey ellipses) (chlorophyll autofluorescence/side scatter) in a clonal culture of 2n-flagellated cells. (c) Removal of reverted cells in 2n-flagellated cultures following EhV infections (120 h post-infection). The fraction (%) of 2n-flagellated and reverted cells is indicated (n = 3). (d) EhV production in 2n-flagellated cultures with reverted cells (n = 3). In c and d, non-infected controls were used for comparison.

### Life cycle reversal: detection of calcified cells in 2n-flagellated cultures

A total of eleven 2n-flagellated (decoupled) strains (LC4-12A-G and LC5-12A-D) were isolated during EhV-infections of RCC1216. Between 9 to 12 months of isolation, calcified cells were detected by microscopy in all 2n-flagellated cultures. As example, in LC4-12A, the fraction of calcified cells accounted for 21% of the total cells, as determined by flow cytometry (Fig. 1b, c). The new calcified cells are referred as reverted cells. Subsequent addition of the virus EhV 201 to all 2n- flagellated cultures lead to the selective lysis of reverted cells, but not 2n-flagellated cells and resulted in viral production (Fig. 1b, c and d). The decoupled strain LC4- 12A was randomly used in ensuing detailed 3D-microscopy and transcriptome analyses to further examine the characteristics of the 2n-flagellated life phase.

### Cell and nuclear ultrastructural differences between cell types

Cellular ultrastructures of 2n-calcified, 1n-flagellated, 2n-flagellated and reverted cells were analysed by FIB-SEM. Cells from each life phase were segmented and volumetrics of organelles (chloroplast, mitochondria and nuclear compartments, such as electron-dense chromatin and nucleolus) were measured. (Fig. 2a).

**Figure 2.**
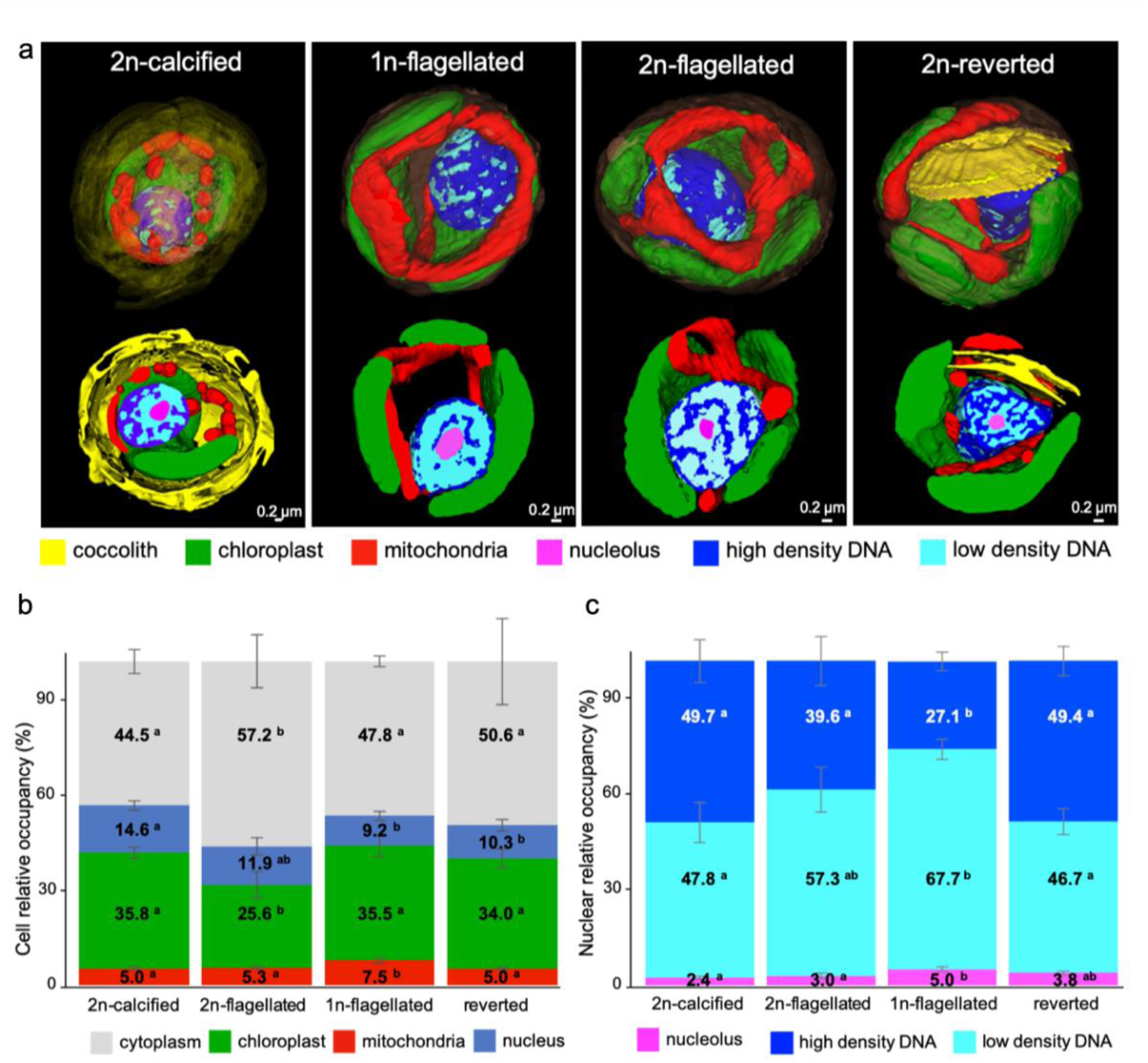
Ultrastructural differentiation of the different life stages of *Gephyrocapsa huxleyi*. (a) 3D reconstitution with FIB-SEM (focused ion beam scanning electron microscopy) of 2n-calcified, 1n-flagellated, 2n-flagellated (decoupled) and reverted (decoupled calcifying) cells. Chloroplasts, mitochondria, nuclear areas and coccoliths are painted in different colors. The scale bar represents 2µm. 1n and 2n: haploid and diploid respectively (b) Relative proportions (% of cell volume) of cytoplasm, chloroplast, mitochondria and nucleus in the four cell types, were calculated from the 3D reconstructions. Significant differences (p < 0.05) identified by Tukey HSD tests are indicated with different letters above the bars (a,b, ab).(c) Relative proportions (% of nuclear occupancy) of high-density and low-density areas of the nucleus and the nucleolus in the four cell types. Significant differences (p < 0.05) identified by Tukey HSD tests are indicated with different letters above the bars (a,b, ab). Statistical analysis was performed using ANOVA type II and Tukey HSD tests to identify significant differences across conditions for each organelle type. Measurements were based on five 2n-calcified cells, five 1n-flagellated cells, six 2n-flagellated cells, and four reverted cells.

FIB-SEM results showed that cell volume was higher in 1n-flagellated and 2n- flagellated (data not shown). Strikingly, reverted cells were 3-fold more voluminous than 2n-calcified cells and coccolith formation was observed in the vacuole in some cells (fig.2a). In 2n-calcified and 1n-flagellated cells, chloroplasts occupied a similar cellular volume (35.8% ± 1.7% and 35.5% ± 3.4% of the cell, respectively). Despite their higher cellular volume, chloroplasts in reverted cells accounted for similar proportions (34.0% ± 2.9%). By contrast, in 2n-flagellated cells chloroplast only represented 25.6% ± 4.1% of the cellular volume. Mitochondria were largest in 1n- flagellated cells, occupying 7.5% ± 0.8% of the cell volume, compared to about 5 % in other cell types (2n-flagellated, reverted and 2n-clacified cells).

The nucleus was proportionally larger in 2n-calcified cells (14.6% ± 1.5% of cell volume) compared to 11.9% ± 2.7% in 2n-flagellated cells, 10.3% ± 1.8% in reverted cells and 9.2% ± 1.4% in 1n cells (fig. 2b). Within the nucleus, variations in electron density patterns were detected, namely high and low density DNA (chromatin) and nucleolus (Fig. 2 a, c). We therefore assessed the volume occupancy of these nuclear compartments (% of the nuclear volume) (fig. 2c). The 2n-calcified and reverted cells exhibited similar density patterns, with an average occupancy of ∼49% for high-density chromatin, ∼47% of low-density chromatin and between 2.4% to 3% for the nucleolus. In contrast, haploid cells (1n-flagellated) displayed an average occupancy of 27% for high-density chromatin (two-fold less than 2n calcified cells), nearly 68% for low-density chromatin and about 5% for nucleolus. Finally, 2n- flagellated cells displayed an intermediate profile, with average occupancies of nearly 40% for high-density chromatin, ∼57% for low-density chromatin and 3% for the nucleolus.

### Cell growth parameters and global transcriptome features

Gene diversity and expression differences between life phases were examined by transcriptome analyses. We note that transcriptome analyses were undertaken prior to the production of reverted cells in LC4-12A, thus solely 2n-calcified, 1n-flagellated and 2n-flagellated cells are included. Inspection by light microscopy validated that each culture contained only one cell type. Cells were grown in replete medium and harvested during exponential growth (Fig. S1b and Table S1). Variations in growth rates were detected one day prior to sampling for the transcriptomes. The 2n- calcified cells displayed higher growth rates (1.33 ± 0.04 day^-1^), followed by 1n- flagellated cells (0.72 ± 0.02 day^-1^) and then 2n-flagellated cell with lower growth rate dynamics (0.47 ± 0.04 day^-1^). All cultures maintained high photosynthetic quantum yield of photosystem II (Fv/Fm) throughout cell growth. At the time of biomass harvesting, Fv/Fm equalled ∼ 0.80 in all strains (Fig. S1c).

To evaluate transcriptional profiles, we defined as transcriptome richness the ratio of expressed transcripts to the total number of transcripts within this ‘synthetic life phase transcriptome’. It is important to note that we set a conservative expression threshold (> 0.01 TPM) to define a transcript as expressed or not. Therefore, we do not rule out the presence, in this core of transcripts, of sequences presenting only residual expression levels. Transcripts with less than 3 raw counts across the 9 libraries were filtered-out, yielding 27350 expressed transcripts. Among these, an average of 23307 (85.2%) were expressed in the 2n-calcified cells, 25263 (92.4%) in the 1n-flagellated cells and 25543 (93.4%) in the 2n-flagellated ones (Fig. 3a), aligning with the observed nuclear patterns. Irrespective of expression magnitude or category-specific differential expression, >78% (21434) of gene-transcripts were common to all three life phases (Fig. 3b). A larger overlap was detected between flagellated cell types (1n-flagellated and 2n-flagellated,11.6%, 3176) than between either of the diploid (2n-calcified and 2n-flagellated ;2.7%, 750) or the non-decoupled stages (1n-flagellated and 2n-calcified, 1.3%, 344). Only a small subset of transcripts was unique to each phase: 1.6% (431) in 1n-flagellated, 3.3% (910) in 2n-calcified cells, and only 1.2% (316) in the 2n-flagellated cells. Independent analyses by RT-qPCR on a set of 19 gene-transcripts validated the specify of expression of the targeted transcripts (Table S2).

**Figure 3.**
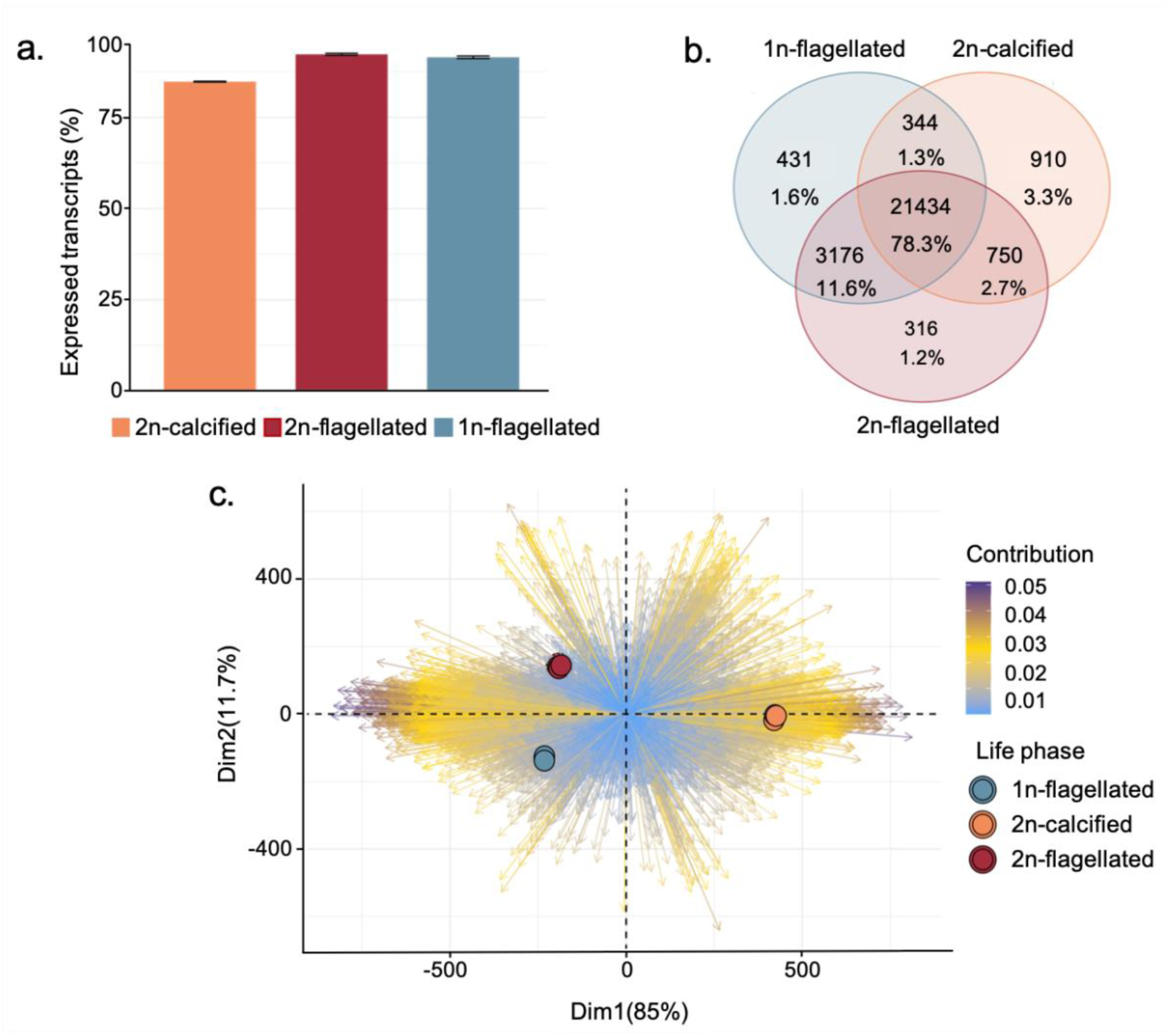
Global properties and differentiation of the transcriptomes of life cycle phases of *Gephyrocapsa huxleyi*. (a) Transcriptome richness of the 2n-calcified, 2n-flagellated and 1n-flagellated. Richness was defined as the ratio of expressed transcripts to the total number of transcripts. A transcript was considered expressed if > 0.01 TPM. (b) Venn diagram of transcript specificity across life phases, in the 1n-flagellated, 2n-calcified and 2n-flagellated. Number of transcripts in each category, and corresponding relative proportions (%) are provided. (c) Principal Component Analysis (PCA) of transcriptional expression profiles and relative separation between different life phases along first 2 principal components (PCs). Each point represents a sample, colored according to its life phase. Arrows represent the contribution of individual transcripts to the PCA, with the color gradient indicating the magnitude of their contribution (blue: low contribution, purple: high contribution). x and y axes: standardized values of the PCs (arbitrary units). 1n and 2n: haploid and diploid respectively

Principal Component Analysis (PCA) delineated the variance between the three life phases (Fig. 3c), with PC1 accounting for 85% of the variation, separating flagellated (haploid and decoupled) from 2n-calcified cells. PC2 encompassed 11.7% of the variation, with 1n-flagellated and 2n-flagellated cells well separated. The top 50 transcripts predominantly influencing PC1 were mostly associated to a morphotype: 34 unique to flagellated forms, two to calcified, nine to diploid, and two commonly detected across all conditions, albeit with expression differing by more than tenfold between flagellated and calcified states (see Table S4). Among these, 34 transcripts annotated with KEGG, KOG or GO terms, were linked to energy metabolism (including nitrogen), amino acid and inorganic metabolism, as well as genetic information processing (transcription machinery, chaperone and folding catalysts, chromosomal proteins, and the ubiquitin system). In contrast, transcripts central to PC2 demonstrated different specificities: two associated with flagellated morphotypes, 19 with diploid-specific expressions, 13 with decoupled cells and ten as core transcripts with significant differential gene expression (DGE) across conditions. Additionally, two transcripts were exclusive to the haploid phase and four were associated with non-decoupled phases (Table S4). These transcripts were related to genetic information processing, including transcription, RNA processing and modification, replication, recombination, and post-transcriptional modifications, as well as lipid biosynthesis and photosynthesis.

### Transcriptional differentiation and functional enrichments

We identified transcriptional differences between 2n-calcified, 1n-flagellated and 2n- flagellated cells. Abundance estimations were performed at the gene level and 23048 transcripts (84.3%) were statistically differentially expressed (False Discovery Rate (FDR) < 0.05). Of these, 9114 transcripts exhibited at least a fourfold change in expression in one pairwise comparison (log2 fold-change) (Table S5). Cluster analysis with k-means resulted in 10 clusters with unique patterns of expression (Fig. 4, Table S6) examined for functional enrichments (GO, KEGG and KOG). In order to reduce terms redundancy of the presented Biological Processes (GO- BP, Fig. 4), significantly enriched terms were grouped under their direct GO ancestor term. Refer to Table S7 for complete results concerning GO-molecular functions, KEGG and KOG. Below are described the enrichments identified in each cluster across all annotation systems.

**Figure 4.**
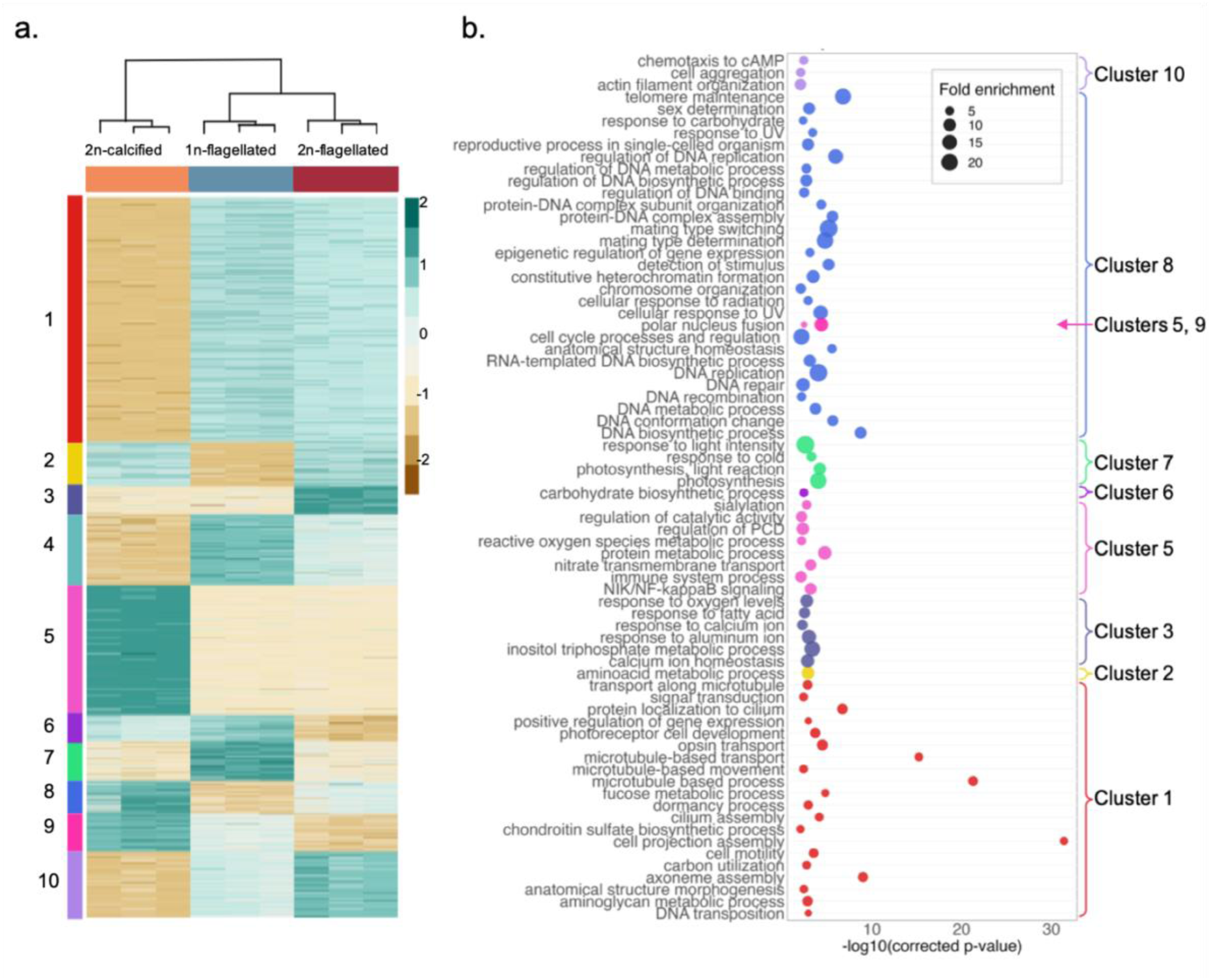
Expression profiles of transcripts in the life phases of *Gephyrocapsa huxleyi*. (a) K-mean (km=10) clustering of transcripts with significant variations of expression (log2 fold change > |2|) between life phases. Expression is standardized by transcript (z-scale). Dark green, high expression levels; brown, low expression levels. 1n and 2n: haploid and diploid respectively. b. Most significantly enriched GO terms (Biological Processes), (hypergeometric test, adjusted p value < 0.05) in the clusters presented in a. Terms were grouped under parental term to reduce redundancy and facilitate visualization. For a complete list of enriched GO terms in each cluster, see Supporting information table S8.

Clusters 1 and 5 accounted for over half (52%) of the differentially expressed genes and contained the most influential transcripts in the PCA displayed in Fig. 3c (Tables S4, S6). These clusters exhibited expression patterns specific to morphotypes (calcified vs non-calcified, flagellated cell types). Cluster 1 (3097 transcripts) was overexpressed in flagellated forms (1n and 2n), featuring enrichments in cytoskeleton and flagellar processes (cilium or flagellum dependent cell motility, microtubule and cilium formation, microtubule-based transport, dynein and catenin binding), as well as transport and metabolism pathways such as carbohydrates (glycosaminoglycan, fucose), amino-acids, nucleotides and DNA mediated transposition, as well as DMSP-dethiomethylase activity (Table S7). By contrast, cluster 5 (1644 transcripts) with transcripts overexpressed in calcified cells, featured enrichments for inorganic ions transport and metabolism (NO_3_^-^, NO_2_^-^), as well as RNA transcription, processing and modification and cell cycle control and chromosome partitioning (Table S7). Necrotic cell death, regulation of reactive oxygen species as well as sialic acid transferase were also associated to cluster 5.

Clusters 4 (900 transcripts) and 10 (823 transcripts) represented transcripts overexpressed in flagellated cells (1n and 2n). Cluster 4 was associated with coenzyme and nucleotide transport and metabolism (Table S7). Cluster 10 (with higher expression in 2n-flagellated than in 1n-flagellated cells) was enriched in cytoskeleton and cellular architecture organization (actin filament, calmodulin binding), in lipid and secondary metabolism associated to energy production and conversion (glucose metabolism), calcium transmembrane transport as well as RNA processing, modifications and chromatin structure and dynamic (Table S7).

Clusters 2 (569 transcripts) and 8 (414 transcripts) comprised transcripts overexpressed in 2n-calcified and 2n-flagellated. Cluster 2 was associated with eukaryotic clusters of orthologous proteins (KOGs) related to transcription, processing and modification of RNA and signal transduction mechanisms (Table S7). Cluster 8 (414 transcripts), with more elevated expression in the 2n-calcified phase, was enriched in the DNA-related functions replication, recombination and repair (DNA replication, post replication repair), as well as control of the cell cycle, cell division and chromosome partitioning (including control of mitotic cell cycle, nuclear chromosome segregation and sister chromatin cohesion). Processes related to the structure of the chromatin and its dynamic were also enriched in cluster 8 (chromatin binding, DNA conformation change, chromosome condensation) as well as mating type determination, switching and reproductive process in single-cell organism) and response to various stimuli (UV, carbohydrates).

Clusters 6 (337 transcripts) and 9 (481 transcripts), represented non-decoupled cell transcripts, overexpressed in both 1n-flagellated and 2n-calcified cells. Cluster 6 was associated with carbohydrate biosynthesis and organic cyclic compound binding. It was additionally associated to intracellular trafficking and DNA replication, recombination, repair and post-translational modifications (Table S7). Cluster 9 (more prevalent in 2n-calcified cells) displayed enrichments in carbohydrate metabolism, polar nucleus fusion and cinnamyl and coniferyl-alcohol dehydrogenases activity (Table S7).

Cluster 3 (353 transcripts) was the only group specific to 2n-flagellated (decoupled) cells. It was associated to transcription, replication, recombination, repair and chromatin structure (Table S7), as well as phosphatidyl inositol binding, response and regulation to calcium ion and aluminum.

Finally, cluster 7 (496 transcripts) was haploid specific and included enrichments in photosynthesis-related functions (chlorophyll and pigment binding) and processes, as well as signal transduction, energy production and conversion, carbohydrate and lipid metabolism and intracellular trafficking and vesicular transport.

### Expression of histone, chromatin and transcription factors

FIB-SEM observations, highlighting marked differences in nuclear ultrastructure between life cycle phases, stimulated the bioinformatic examination of genetic functions potentially involved in chromatin remodeling and likely implicated in life cycling. Alongside, several functional enrichments, particularly those related to general DNA replication and repair, chromosome conformation and regulation of gene expression, underscored the potential role of epigenetic regulation in differentiating life phases. Thus, we zoomed on transcripts associated with the general chromatin machinery: histones, readers, writers, remodelers, chaperones, as well as transcription factors (TFs) (Fig. 5). We clustered through k-mean analysis those that were associated to a log2 fold-change > |0.5|.

**Figure 5.**
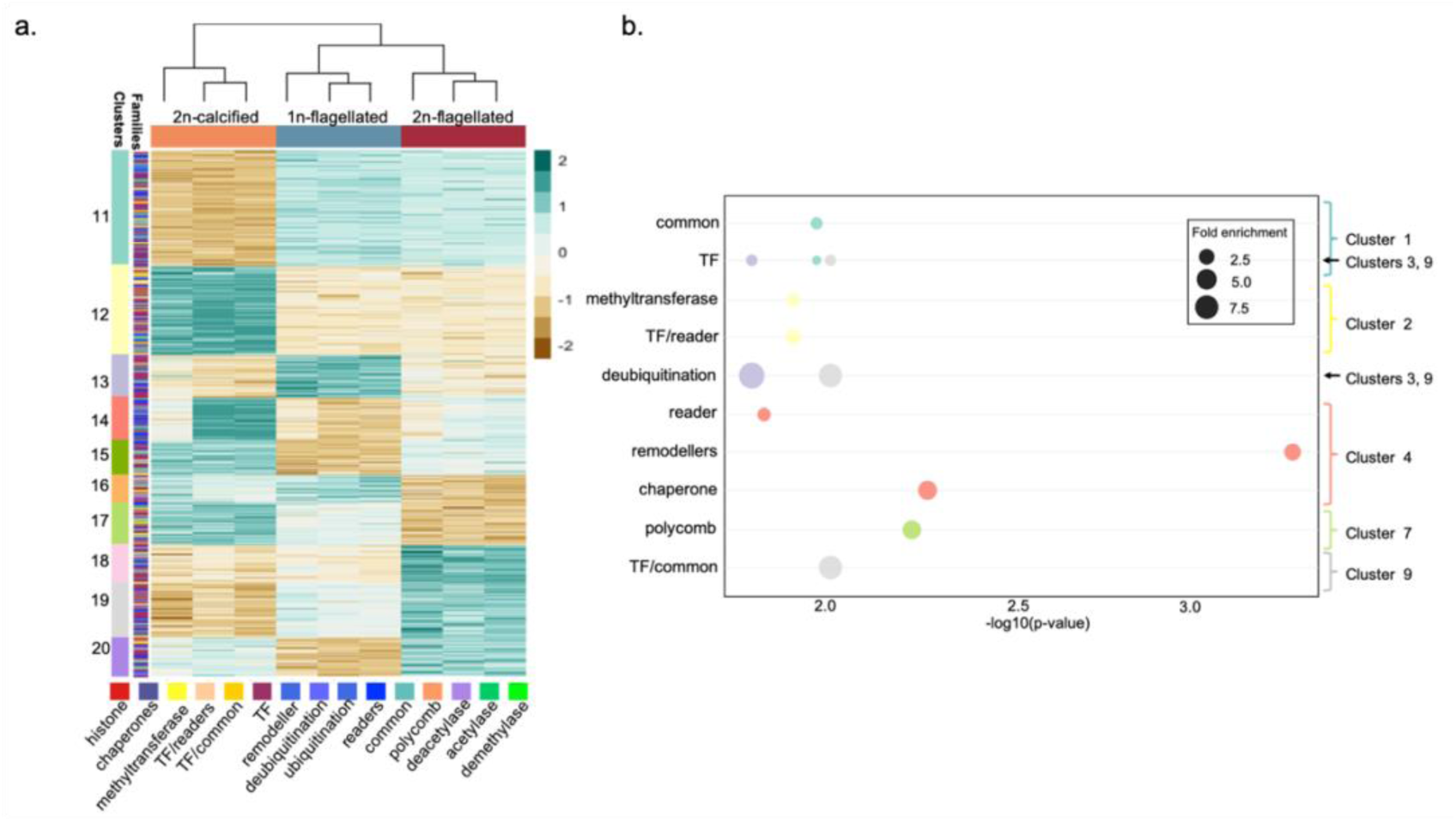
Expression profiles of chromatin associated transcripts related to epigenetic functions in the life phases of *Gephyrocapsa huxleyi*. (a) K-mean (Km=10) clustering of transcripts associated to statistically significant variations of expression (FDR < 0.05, log2fold change >|0.5|) between life phases. Expression is standardized by transcript (z-scale). Dark green, high expression levels; brown, low expression levels. 1n and 2n: haploid and diploid respectively (b) Significantly enriched epigenetic families, (hypergeometric test, adjusted p value < 0.05) in clusters presented in (a). “common”: transcripts for which only one protein domain, commonly found to be associated with chromatin-related proteins and processes, could be identified. In the lack of other identifications, transcripts family could not be determined (conserved domains: Dpy-30, Kelch, WD-40, zf-C2H2 and zf-CXXC).

Ten clusters were defined, each with highly heterogeneous gene family composition. They are numerated from 11 to 20 to avoid potential confusion with the above mentioned results. Clusters 11 and 12 contained more than a third of the transcripts, segregating cells by their morphotypes. Cluster 11 (146 transcripts), primarily expressed in flagellated cells, was enriched in TFs and protein domains commonly associated with the chromatin machinery (‘common’), though no specific family was identified. Cluster 12 (111 transcripts), highly expressed in calcified cells, was enriched in chromatin specific methyltransferases, TFs, and chromatin readers.

The other clusters were considerably smaller and associated to specific life phases. Cluster 19 (69 transcripts), expressed in both flagellated conditions, was more abundant in the 2n-decoupled cells compared to the 1n-calcified cells. This cluster was associated with TFs, proteins involved in deubiquitination processes, and ’common’ domains. Clusters 14, 15, and 20 (54, 43, and 47 transcripts, respectively) were expressed in diploid cells. Cluster 14, characterized by clear heterogeneity among its replicates and higher expression in diploids, was enriched in readers, chaperones, and remodelers. Clusters 15 and 20, more abundant in the 2n-calcified and 2n-flagellated cells respectively, had no significant enrichments. Cluster 13 (53 transcripts) was the only group characterized by an overexpression exclusive of the 1n-flagellated cells and was associated with TFs and deubiquitination proteins.

Cluster 18 (47 transcripts), the only group overexpressed in the decoupled cells, was not associated with any significant enrichments. Finally, clusters 16 and 17 (35 and 53 transcripts, respectively) were more expressed in non-decoupled cells. Cluster 16, with higher expression in 1n-flagellated cells, was not associated with significant enrichments. Cluster 17, with transcripts more expressed in the 2n-calcified than in the 1n-flagellated cells, was associated with the Polycomb protein complex.

Twenty-two transcripts were associated to canonical histones or their variants in our dataset: 10 histones H2A, 2 H2B, 11 histones H3 and 1 H4. Four transcripts were associated to the histone family without a clear identification (Table S5). Five histones H2A were highly expressed in all cells and 2 of them were strongly depleted in the 2n-flagellated and 1n-flagellated respectively. One H2B was over 50% more expressed in 2n-calcified cells. Four histones H3 were abundant and strongly overexpressed in one life phase. One was expressed only in the haploid cells and 1 was expressed in the diploid phases and more abundant in the 2n-flagellated. Two other histones H3 were overexpressed in the 2n-calicified cells compared to the 2 other conditions. (Table S5).

One TALE-HD TF was identified and expressed only in the haploid cells. This specificity was validated by RT-qPCR (Table S2).

### Flagellated cell-specific expression during viral infection

Earlier work showed that decoupled cells are produced during Ehv infection (Frada et al. 2017). To test life cycle transition during viral infection and gain insight on mechanisms of 2n-flagellated production, we re-examined published transcriptomes of the diploid *G. huxleyi* strain CCMP2090 interacting with EhV201 (1 and 24 hr post infection) (Rosenwasser *et al*. 2014). We targeted transcripts that demonstrated a high homology level (see material and methods) with flagellated-specific transcripts identified in our work.

We identified 176 gene homologs in this dataset. Through k-means clustering, four co-expressed groups were identified and functional enrichments were examined (Fig. 6a; Table S3 for complete list of clustered transcripts, associated annotations, Table S7 for enrichments). Clusters 1 and 2 grouped transcripts with higher expression levels in control (non-infected) condition. Cluster 1 (13 transcripts) was found to be enriched in organelle organisation process and in general metabolism annotations, as well as with protein families involved in genetic information processing (mRNA biogenesis, chaperones and folding catalysts). Cluster 2 (22 transcripts) was enriched in general metabolism pathways (carbohydrate, sugar, lipid). Its transcripts were also related to proteins involved in genetic information processing (transcription and translation machinery, chromatin structure). Cluster 3 (22 transcripts) comprised transcripts that were upregulated at 1h post-infection but downregulated by 24h. This cluster was enriched in protein modification process, and its transcripts associated to the metabolism of amino acids, the ubiquitin system and cilium and its associated proteins. Cluster 4, containing 68% (119 transcripts) of all clustered transcripts was markedly upregulated 24h post-infection. It was enriched in orthologous groups and functions associated to the cell cycle control and division and partitioning of chromosomes, to signal transduction and cytoskeleton mechanisms (tubulin binding, microtubule cytoskeleton organisation), as well as post-translational modifications, chaperones and protein turnover (histone methyltransferase activity, nucleotide/nucleoside biosynthetic processes and binding) and intracellular trafficking and carbohydrate transport and metabolism.

**Figure 6.**
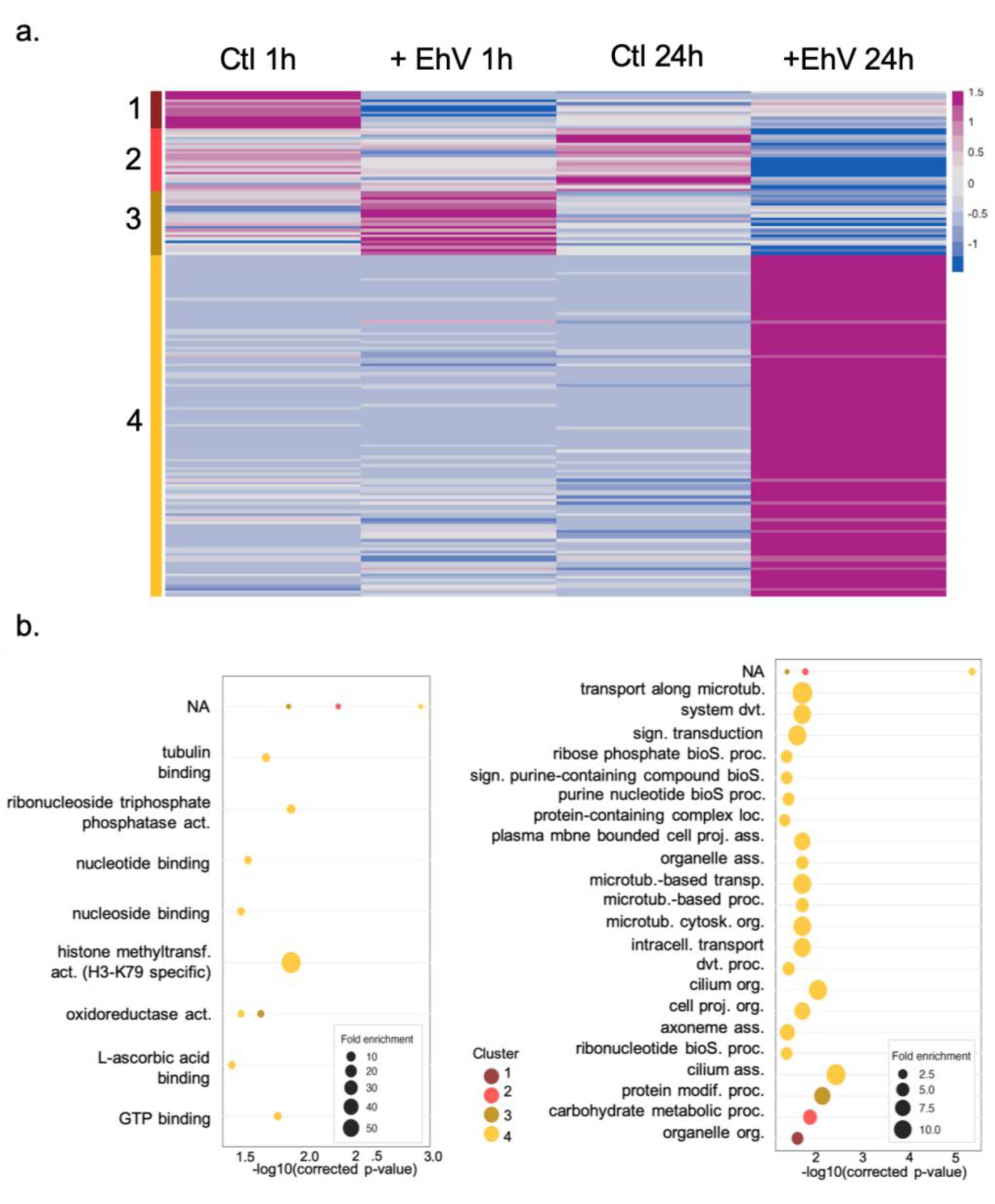
Expression profiles of “flagellated-specific” genes in the *Gephyrocapsa huxleyi* strain CCMP2090 during viral infection by EhV201 and non-infected control. (a). K-means clustering (Km=4) of homologs in CCMP2090 to flagellated transcripts in absence (control, ctl) or presence (+EhV) of viral infection, 1h and 24h post infection. Expression is standardized by transcript (z-scale). This heatmap uses different colours relative to fig. 4 and 5 to emphasize the different dataset represented here (CCMP2090 vs life phase transcriptome for fig 4 and 5, resp.). Dark pink; high expression level, Blue, low expression level. (b) Significantly enriched GO terms (left panel -molecular functions, right panel-biological processes), (hypergeometric test, adjusted p value < 0.05) in cluster presented in (a). abbreviations: microtub.: microtubule, dvt.: development, sign.: signal, bioS.: biosynthetic, proc.: processes, loc.: localization, mbne.: membrane, proj.: projection, ass.: assembly, transp.: transport, cytosk.: cytoskeleton, org.: organization, intracell.: intracellular, modif.: modification

## Discussion

The life cycle, encompassing all cellular stages, is a fundamental attribute of eukaryotes (Mable & Otto, 1998), underpinning their range of biological and ecological functions and their capacity to adapt to environmental changes through cell differentiation. In marine phytoplankton, the understanding of life cycle versatility and regulation remains limited. Here, we examined the life cycle of the cosmopolitan, bloom-forming coccolithophore *G. huxleyi*, which includes 2n-calcified cells, 1n- flagellated cells and 2n-flagellated cells produced during viral infections (Frada *et al*., 2017). We specifically investigated the fate of the virus-resistant 2n-flagellated cells during culture observations and the morphogenetic differentiation between 2n- calcified, 1n-flagellated and 2n-flagellated cell types through ultrastructural and transcriptome analyses.

A key finding of this study is the ability of 2n-flagellated cells to revert back to a calcifying morphotype in the absence of viral pressure. Given the low proportion of newly calcifying cells (reverted) within the 2n-flagellated cultures (20%), it is reasonable to assume that only a small subset of cells undergoes flagellate-to- calcified switch. Reverted cells subsequently proliferate mitotically, alongside flagellated cells, becoming detectable by microscopy perhaps only weeks post-transition.

Morphological transformations into a resistant phenotype in response to viral infections, followed by a reversal to the original cell phenotype, have been observed in other protists. For example, certain diatom and raphidophyte species can produce viral-resistant cysts during infections, which germinate back to the vegetative cell-type in the absence of viruses (Dingman, 2015; Pelusi *et al*., 2021). However, the role of viral infection in triggering life cycle transitions and the subsequent ability to reverse this process, has been to our knowledge uniquely described here, in *G. huxleyi*. The production of decoupled cells resembling apospory observed in algae and plants (e.g., Cock *et al*., 2014) is thought to represent a natural escape strategy from EhV infection (Frada *et al*., 2008, 2017). However, the reversal to a calcifying state suggests a potential cost for diploid cells in maintaining a haploid-like phenotype. The cost of viral resistance is a common phenomenon in bacteria and protists (e.g, Lenski & Levin, 1985; Tomaru *et al*., 2009; Thomas *et al*., 2011; Dingman, 2015). In *G. huxleyi*, as corroborated by a previous study (Frada *et al*., 2017), 2n-flagellated (decoupled) cells display reduced growth rate and carrying capacity compared to both 2n-calcified and 1n-flagellated (haploid) cells. This may be explained by their inherently larger cell volume, revealed here with electron microscopy, and ensuing higher nutrient requirements for cell division. In addition, FIB-SEM measurements indicate a relative proportion of chloroplasts ∼25% lower in the 2n-flagellated cells compared to other cell types, suggesting a reduced capacity to generate energy, while the mitochondrial volume in these cells, does not change. Thus, the phenotype reversion (2n-flagellated to 2n-calcified) likely restores the metabolic and physiological advantages.

We also observed that the reversion of 2n-flagellated cell in *G. huxleyi*, is associated with a loss of the viral resistance. Calcified reverted cells, once re-exposed to EhV, were eliminated from the cultures. Mechanisms underlying *G. huxleyi*’s sensitivity and resistance to EhV may involve a variety of metabolic processes, reflecting the complex host-virus ‘arms-race’ dynamics (Bidle & Vardi, 2011; Bidle, 2015). Previous work linked specific sphingolipid molecules (sialic acid sphingolipids, sGSL) to EhV sensitivity in diploid cells (Fulton *et al*., 2014). sGSLs were proposed to play an intrinsic role in coccolith biosynthesis and serve as ligands for viral attachment and entry. These sGSLs are not biochemically detected in haploid cells, which are insensitive to EhV (Hunter *et al*., 2015). Enrichment for sialylation transcripts (cluster 5, Fig. 4) was observed only in diploid-calcified cells and not in flagellated cell types (Table S7). This suggests that sGSLs are likely absent in 2n-flagellated cells but may reappear in reverted cells, potentially restoring viral sensitivity.

Ultrastructural analyses enabled the quantitative assessment of significant differences in the nuclear architecture between cell-types. Notably, nuclei of 2n- calcified and reverted cells displayed nearly equal parts of low- and high-density chromatin, accounting for 47-49% of the nuclear volume, respectively. In contrast, nuclei of 1n-flagellated were predominantly of low-density chromatin (> 67%), while the nuclei of 2n-flagellated cells displayed an intermediate profile (Fig. 2c). Specific approaches are needed to confirm the nature of high- and low-density areas (‘Chromatin accessibility profiling methods’, 2021). However, it is reasonable to assume that they correspond to heterochromatin (densely packed and usually transcriptionally inactive), and euchromatin (relaxed and generally associated with a more active transcriptional state of the DNA), respectively (Huisinga *et al*., 2006). In eukaryotes, chromatin packaging is highly dynamic and central for spatiotemporal regulation of transcription in many cellular processes, including cell differentiation, development and life cycle in animals, plants and algae (Zheng & Xie, 2019; Vigneau & Borg, 2021). Rearrangements in chromatin ultrastructure is often mediated by extensive epigenetic reprogramming, involving hPTMs and the activity of TFs that modulate transcriptional landscapes (Morgan *et al*., 2005; Thiriet-Rupert *et al*., 2016; Vigneau & Borg, 2021). Thus, the observed nuclear ultrastructural patterns, across the different phases of the life cycle in *G. huxleyi,* through unprecedented volume quantification, suggest vast divergence of gene expression between cell types, with 2n-flagellated cells maybe displaying an intermediary transcriptional profile.

Complementary comparative transcriptome analyses enabled to test this hypothesis, while gaining insights on wider functional attributes of 2n-calcified, 1n-flagellated and 2n-flagellated cells during exponential growth. Conservative identification of common and phase specific transcripts, highlighted that the strong majority of transcripts were shared between the three life phases (>78%), regardless of expression levels (Fig. 3b). A more important overlap was observed between flagellated morphotypes (11.6%) compared to the diploid cells (2.7%) (Fig. 3b). Moreover, as revealed by the PCA (Fig.3c), the main factor of separation between samples (>85% of the variance) was attributed to the cell morphology (flagellated vs calcified), while only 11.7% of the variance equally separated 1n and 2n-flagellated from 2n-calcified (Fig. 3c). This separation might lay in the mechanisms of production of these cells, haploid cells being produced by meiosis while decoupled cells might originate from apospory-like mechanisms. Thus, from a transcriptomic perspective, 2n-flagellate cells do not represent an intermediary level of expression but are in fact much closer to the haploid profile. Regulation of cell morphology, rather than the ploidy level itself, is the main factor of differentiation between cell types in *G. huxleyi*, as similarly detected in brown algae, displaying a haplodiplontic life cycle and apomictic variants (Arun *et al*., 2019). Examination of DGE and enrichment profiles further highlighted the contrasts between flagellated and 2n-calcified cells. Namely, enrichments in functions related to cell motility were identified in flagellated cell types. By contrast, enrichments in functions related to the transport and metabolism of inorganic ions were associated to 2n-calcified cells. This can be associated with flagella biosynthesis, signalling and with calcification, respectively, in broad agreement with previous transcriptome comparisons of 1n-flagellated and 2n-calcified cells (von Dassow *et al*., 2009; Rokitta *et al*., 2011).

Significant differences between cell types were also detected in functions associated to chromatin remodelling and transcriptional regulation (including transcription factors, TFs). At this level, morphotype is still the main differentiating factor of transcript expression (Fig.5). A majority of transcripts were either over-expressed in the calcified cells or in the flagellated cells, further emphasizing phenotype as the main manifestation of gene regulation in *G. huxleyi* life phases, independently of ploidy levels. We detected for example an over expression in the 2n-calcified cells, associated to histones methyltransferases (Fig. 5a, cluster 12). It can align with the high chromatin compaction (Chen & Dent, 2014) detected in these cells (Fig. 2b).

Additionally, ubiquitin removal (deubiquitination) enrichments were associated to transcripts with clear DE between cell types, a function associated with chromatin relaxation and increased transcriptional activity (Mikulski *et al*., 2017). It fits overall patterns of chromatin, less dense in flagellated cells compared to 2n-calcified cells (Fig. 5a, clusters 13 and 19). Enrichments observed in cluster 14 (Fig. 5a) were also relevant. We observed a heterogeneous gradient of expression from 2n-calcified to 2n-flagellated cells, in transcripts associated to chromatin "architects" (remodelers, readers and chaperones) (Clapier & Cairns, 2009). suggesting intermediate level of expression and recruitment. We think it might further be associated with a transitional level of chromatin organization between the two types of diploid cells. We also identified enrichments for the Polycomb-Complex, associated with transcripts over-expressed in 2n-calcified cells compared to 1n-flagellated cells; a TALE-homeodomain TF specific to 1n cells and histone H3 variants with marked DGE between life phases. All these gene-functions are conserved across eukaryotes and play central roles in the regulation of life cycle transitions (Horst *et al*., 2016; Mikami *et al*., 2019; Dierschke *et al*., 2021; Lee *et al*.). Polycomb associated transcripts in our dataset, likely correspond to the Polycomb Repressive Complex 2 (PRC2, Grau-Bové *et al*., 2022). PRC2 mediates the trimethylation of histones 3 (H3) on lysine 27 (H3K27me3), associated to the silencing of genes encoding TALE-HD TF (KNOX and BELL). These transcription factors are crucial for haploid-to-diploid transitions in plants and algae (Yelagandula *et al*., 2014; Wollmann *et al*., 2017; Thangavel & Nayar, 2018; Loppin & Berger, 2020; Vigneau & Borg, 2021; Hirooka *et al*., 2022) (Loppin & Berger 2020, Wollman et al. 2017, Yelagandula et al. 2014, Vigneau and Borg, 2021). Overall, these results suggest the involvement of conserved key mechanisms in the life cycle differentiation in *G. huxleyi*.

Finally, we screened for homologs of flagellated-specific transcripts (expressed only in the 1n- and 2n-flagelled cells) in *G. huxleyi* strain CCMP2090, during EhV infection (Rosenwasser *et al*., 2014). A variety of transcripts mainly involved in metabolic functions were detected in control (non-infected) conditions, indicating that life cycle specificity of various functional genes can change between *G. huxleyi* strains and likely as well between growth conditions and external environmental stimuli. However, most of the flagellated-cell homologs identified were over- expressed 24h post-infection, strongly supporting life cycle transition during EhV infections (Frada *et al*., 2017). Among flagellated-cell homologs, we detected a wide set of motility and cell architecture related genes, as well as signaling transduction mechanisms. We note that CCMP2090 does not form flagella, albeit a strong up- regulation of some flagellar functions upon infection (Frada *et al*., 2017). This morphological impairment likely results from the genome erosion of this strain and subsequent loss of functional flagella biosynthesis genes (von Dassow *et al*., 2015). In any case, CCMP2090 can form viral resistant cells that display organic-body scales, only featured in flagellated cells (Frada *et al*., 2017). We further detected the association of these flagellated specific transcripts with functions involved in gene expression, including a strong enrichment in H3K79 specific methyltransferase that regulates diverse cellular processes, such as development, reprogramming, differentiation and proliferation (Feng *et al*., 2010; Takahashi *et al*., 2011; Tatum & Li, 2011). It is generally associated to active transcription and maintains chromatin accessibility for histone acetylation and transcription factor binding (Godfrey *et al*., 2019). Although limited, these lines of evidence further recall the involvement of the upstream epigenetic regulation during the differentiation of 2n-flagellated cells in *G. huxleyi* during viral infection.

In conclusion, our study provides detailed comparative assessment of three life-cycle cell types in *G. huxleyi*, evidences for life cycle versatility during viral interactions and novel insight on chromatin remodeling and transcription regulators, which likely play a central role in mediating life cycle transitions. These results should provide the basis for further development of future studies on the role of epigenetic control during cell differentiation and life cycle in coccolithophores. Finally, one of the greatest gaps limiting comprehensive understanding about *G. huxleyi* is the lack of knowledge of the occurrence, distribution and role of flagellated cells in natural populations and during blooms where viral infections are recurrent. The analyses presented here, enable to refine useful gene markers for the three life phases of *G. huxleyi* that can be used to interrogate coccolithophore assemblages in the oceans.

## Supporting information

Supporting_informationS1

Supporting_informationS2

## Acknowledgements

This study was supported by the Israel Science Foundation (2921/20) attributed to MJF. JD was supported by CNRS and ATIP-Avenir program funding. We thank Prof. Aharon Kaplan for hosting SF and Prof. Assaf Vardi to enable access to his lab for molecular biology work. We thank The Life Sciences Core Facilities from The Weizmann Institute of Science for the preparation and transcriptome sequencing as well as Esther Feldmesser for support and feedback on bioinformatic analyses. We thank Guy Schoehn and Christine Moriscot, and the electron microscope facility at IBS, which is supported by the Rhône-Alpes Region, the Fondation Recherche Medicale (FRM), the fonds FEDER, the Center National de la Recherche Scientifique (CNRS), the CEA, the University of Grenoble, EMBL, and the GIS Infrastructures en Biologie Sante et Agronomie (IBISA) CK was supported by Academia Sinica (AS-CDA-110-L01) and NSTC, Taiwan (111-2611-M001-008-MY3).

## Author contributions

Conceptualization: MJF, LB, SF; Methodology: MJF, JD, LB, SF, OM; Investigation: LB, SF, YA,GK, FC, BG, RT, YS, MJF; Formal Analysis: LB, YD, SF; Resources: CK, FC, MJF; Data curation: LB, JD, BG, RT, YS; Visualization: LB, JD, RT, YS; Writing (original draft): LB, MJF; Writing (review & editing): LB, MJF, SF, JD, OM and CK; Supervision: MJF, OM; Funding acquisition: MJF, CK.

## Data availability

Data are available in the main text or supplementary materials. Life phase transcriptome predicted coding sequences (cds) are available in supporting information file S2. Raw data have been deposited in the NCBI under BioProject PRJNA1134391.

## Supporting information tables and file

Table S1: Cell density and maximum photochemical quantum yields of PSII measurements

Table S2: RT-qPCR primers and results

Table S3: Functionnal annotation and clustering information of CCMP2090 homolog to flagellated specific transcripts

Table S4: Top 50 transcripts contributing to the variance between sampled (PC1 and PC2)

Table S5: The life phase transcriptome

Table S6: Clustering informations

Table S7: Significant results of functionnal enrichment tests (hypergeometric test, BH correction)

Supporting information file 1:

Methods S1: Bioinformatic parameters

Methods S2: Life phase transcriptome additional information

Methods S3: Local databases for histones, chromatin-associated transcripts and transcription factors

Methods S4: Transcriptome validations by qRT-PCR

Methods S5: Focus Ion Beam-Scanning Electron Microscopy, extended information

Supporting Figure Fig. S1: Characterisation of cultures for transcriptome analysis

Supporting figure Fig. S2: TALE (Three Amino acid Length Extension) Homeoproteins of plants and algae that regulate life cycle transition

Supporting information file 2: Life phase transcriptome cds (fasta file)

