## Supporting_informationS1 for "Life Cycle and Morphogenetic Differentiation in Heteromorphic Cell Types of a Cosmopolitan Marine Microalga": Supporting_informationS1.docx

^1^Department of Ecology, Evolution and Behavior, Silberman Institute of Life Sciences, The Hebrew University of Jerusalem, Jerusalem, Israel; ^2^The Interuniversity Institute for Marine Sciences in Eilat, Eilat, Israel; ^3^Laboratoire Physiologie Cellulaire et Végétale, Univ. Grenoble Alpes, CNRS, CEA, INRAE, IRIG-DBSCI-LPCV, 38000 Grenoble, France; ^4^Translation Genomics Lab and Medical Genetics Institute, Shaare Zedek Medical Center, 93722, Jerusalem, Israel; ^5^Institute of Plant and Microbial Biology, Academia Sinica, Taipei, Taiwan.

°Current address: Faculty of Biology, Technion – Israel Institute of Technology, Haifa 3200003, Israel.

Author for correspondence:

Miguel J. Frada

The following Supporting Information is available for this article:

**Methods S1** – Bioinformatic parameters

**Methods S2** – Life phase transcriptome additional information

**Methods S3** – Assembly of local databases for histones, chromatin-associated transcripts and transcription factors

**Methods S4** – Transcriptome validations by qRT-PCR

**Methods S5** - Focus Ion Beam-Scanning Electron Microscopy, extended information

**Supporting Figure Fig. S1** Characterisation of cultures for transcriptome analysis

**Supporting figure Fig. S2** TALE (Three Amino acid Length Extension) Homeoproteins of plants and algae that regulate life cycle transitions.

**Methods S1 Bioinformatic parameters**

| Program | options |
| --- | --- |
| STAR version 2.7.10a | --alignEndsType EndToEnd --alignIntronMax 2500 --alignIntronMin 30 |
| StringTie V2.2.1 | -c 2 -m 75 |
| Cuffmerge V2.2.1 | -G -s |
| gffread V0.12.7 | default |
| CD-HIT-EST (V4.8.1) | -c 1 -n 11 -g 1 |
| OmicsBox (free version) | default |
| TransDecoder V5.5.0 | --retain_long_orfs_mode dynamic and --single_best_only |
| InterProScan v5.52-86. | -goterms |
| Diamond V2.0.14.152 | diamond blastp --db <database.dmnd> --out <file.out> --outfmt 6 --query <queries.fasta> --ultra-sensitive -evalue 1e-3 |
| Muscle program V5.1 | -align <input_files.fasta> -output <output.aln> -maxiter 40 |
| HMMER program V3.3.2 | hmmbuild –amino <profile.hmm> <input_file.aln>  hmmscan –tblout <out_file.tbl> <profiles.hmm> <query_transcripts.fasta> > <hmm_results.out> |

**Methods S2**

**Life phase transcriptome additional information**

Sequencing was performed on a NextSeq platform, generating a total of 530500896 single-end 75 bp reads. The average read counts were approximately 59.7 million for the 1n-flagellated , 57.6 million for the 2n-calcified, and 59.6 million for the 2n-flagellated libraries, with a GC content of 66.3%, 66.6%, and 66.3% respectively. After quality trimming and the removal of contaminating sequences, about 97% to 98% of the reads were retained.

Despite the availability of a reference genome and multiple transcriptomes from various strains of *G. huxleyi*, the extensive genome variability within this species complex (Read *et al.*, 2013) prompted us to construct a synthetic life phase-specific transcriptome. This approach was intended to minimize the loss of key transcripts due to genetic divergence from the reference strain. We achieved this by aligning reads from each library to a newly sequenced genome from a haploid strain (RCC1217) derived from RCC1216. The mapping rates for these alignments ranged from 91% to 94% (including uniquely and multimapping reads), and the resulting alignments were assembled into potential transcripts using StringTie (Pertea *et al.*, 2015). These transcripts were then merged into a unique set of 56788 non-redundant transcripts.

To assess the completeness of our life phase transcriptome, we utilized BUSCO (Benchmarking Universal Single Copy Orthologs), searching against the Eukaryote database which includes 255 conserved representative groups (Manni *et al.*, 2021). Despite using short single-end reads, 191 (75%) of the conserved orthologs were found complete, with 115 occurring as single copies and 76 duplicated. We also identified 25 (9.8%) as fragmented and 39 (15.3%) were missing. These results align closely with those from recently assembled genomes and proteomes (Skeffington *et al.*, 2023).

Coding sequences (cds) were predicted for transcripts longer than 30 nt using TransDecoder. Of the 56092 retained transcripts, approximately 77.4% of the coding sequences were found to be complete, 17.4% were partial at the 5' end, 2.3% were partial at the 3' end, and 2.7% were internal fragments. Annotation was performed using EggNOG-mapper and Gene Ontology (GO) through Blast2GO.

To identify major transcriptomic differences between the 3 life phases, we quantified reads abundances at the gene level in each library with RSEM (Li & Dewey, 2011). It resulted in a dataset of 27365 representative transcripts prior any filtering steps. Of these, 7477 (27.3%) were associated with a gene ontology, 7639 (28%) were associated to a KEGG number and 11390 (41.6%) were associated with a KOG identifier (including unknown functions).

**Methods S3**

Local databases for histones, chromatin-associated transcripts and transcription factors

Peptide sequences of chromatin-associated proteins (Acetylases, Chaperones, Polycomb Complex, Methylases, Chromatin Readers, and Remodellers), canonical Histones (H2A, H2B, H3, and H4), primary Histone variants (H2A.Z, macro H2A, and cenH3) and specific taxa Histones (Cryptista, n=32; Haptista, n=50; Rhizaria, n=10; Streptophyta, n=137; Rhodophyta, n=40; Chlorophyta, n=189; Stramenopiles, n=231; Alveolata, n=258) were downloaded from the Grau-Bové et al. 2022 Github repository, resulting in 1136 non-redundant histones and 59262 chromatin-associated protein sequences. Transcription factors from various taxa (*Arabidopsis thaliana* , n=2297; *Bathycoccus prásinos*, n=140; *Chlorella variabilis* NC64A, n=164; *Chlamydomonas reinhardti*i, n=231; *Coccomyxa subellipsoidea*, n=139; *Micromonas pusilla*, n=151; *Micromonas sp*. RCC299, n=154; *Ostreococcus lucimarinus*, n=115; *Ostreococcus sp. RCC809*, n=103; *Ostreococcus tauri*, n=100; *Picochlorum sp.*, n=102; *Volvox carteri* , n=197) were sourced from PlantTFDB (<http://planttfdb.gao-lab.org>) and from Thiriet-Rupert *et al.*, 2016 (*Chlamydomonas reinhardtii* , n=212; *Gephyrocapsa huxleyi* ,n=488; *Nannochloropsis gaditana*, n=102; *Pavlova sp*., n=134; *Porphyridium purpureum*, n=190; *Phaeodactylum tricornutum,* n=202) resulting in a non-redundant database of 5219 sequences. Diamond blastp v2.0.15 was used for similarity searches, with life phase transcripts as the database and protein-specific databases as queries. Hits with less than 30% identity over fewer than 90 aligned residues were discarded. For each gene, only the isoform with the lowest e-value was retained. Protein domain identification involved creating multiple sequence alignments with MUSCLE v5.1 (Edgar, 2004, default parameters), constructing HMM profiles with hmmbuild and identifying domains using hmmscan. Hits with an e-value < 0.005 for both the full sequence and the best single domain were discarded. Transcripts were sorted by e-value, retaining those with the best values; discrepancies were manually reviewed. Only transcripts identified by both Diamond blastp and HMMER were retained.

Identification of *Gephyrocapsa huxleyi* TALE-HD Transcription factor:

TALE (Three Amino acids Loop Extension) Homeodomain sequence was identified by local blast against *G.huxleyi* transcriptome using known sequences as the database.

**Methods S4**

**Transcriptome validations by qRT-PCR**

Total RNA was isolated from replicate samples from each cell type. Reverse transcription of purified RNA to cDNA was performed using the SuperScript IV RT-PCR system (Invitrogen), as detailed by the manufacturer. Quantification of transcript levels was conducted utilizing the Fast SYBR Green Master Mix (Applied Biosystems). Specific primers targeting a variety of life phase specific genes were used (details provided in Table S2). Amplification reactions were undertaken on a StepOne Real-Time PCR System (Thermo Fisher Scientific) as follows: 95°C for 2 minutes, followed by 35 cycles of denaturation at 95°C for 15 seconds, annealing at 60°C for 30 seconds, and extension at 72°C for 30 seconds. The relative expression levels of the genes were calculated by employing the 2-ΔΔCt method, with normalization to diploid calcified control samples with Tubulin as the reference gene.

**Methods S5**

**Focus Ion Beam-Scanning Electron Microscopy, extended information**

For Focus Ion Beam-Scanning Electron Microscopy (FIB-SEM), cell pellets were first subjected to freeze substitution (FS). For that a mixture 2% (w/v) osmium tetroxide and 0.5% (w/v) uranyl acetate in dried acetone was used. The FS machine was programmed as follows: 60-80 hr, at -90°C, heating rate of 2°C hr-1 to -60°C (15hr), 10-12hr at -60°C, heating rate of 2°C h-1 to -30°C (15hr), 10-12 hr, at -30°C. Samples were then washed in acetone four times for 20 min at -30°C and embedded in anhydrous araldite. Without accelerator, a graded resin/acetone (v/v) series was used with each step lasting 2h at increased temperature: 30% resin/acetone bath from -30°C to -10°C, 50% resin/acetone bath from -10°C to 10°C, 70% resin/acetone bath from 10°C to 20°C.

Samples were then placed in 100% resin without accelerator for 8-10 hr and in 100% resin with accelerator (BDMA) for 8 hr at room temperature. The sample was mounted onto the edge of a SEM stub (Agar Scientific) using silver conductive epoxy (CircuitWorks) with the trimmed surfaces facing up and towards the edge of the stub. The sample was gold sputter coated (Quorum Q150RS; 180 s at 30 mA) and placed into the FIB-SEM for acquisition (Crossbeam 540, Carl Zeiss Microscopy GmbH). Atlas3D software (Fibics Inc. and Carl Zeiss Microscopy GmbH) was used to perform sample preparation and 3D acquisitions. First, a 1 µm platinum protective coat (20-30 µm2 depending on ROI) was deposited with a 1.5 nA FIB current. The rough trench was then milled to expose the imaging cross-section with a 15 nA FIB current, followed by a polish at 7 nA. The 3D acquisition milling was done with a 1.5 nA FIB current. For SEM imaging, the beam was operated at 1.5 kV/700 pA in analytic mode using an EsB detector (1.1 kV collector voltage) at a dwell time of 8 µs with no line averaging. For each slice, a thickness of 8 nm was removed, and the SEM images were recorded with a pixel size of 8 nm, providing an isotropic voxel size of 8 × 8 × 8 nm3. From the stack of images, regions of interest containing cells were cropped using the open software Fiji (https://imagej.net/Fiji), followed by image registration (stack alignment), noise reduction, semi-automatic segmentation, 3D reconstruction of cells and morphometric analysis as in (Decelle *et al.*, 2021; Uwizeye *et al.*, 2021; Gallet *et al.*, 2024). Image registration was done by the FIJI plugin ‘Linear Stack Alignment with SIFT’ (Lowe, 2004), then fine-tuned by AMST (Hennies *et al.*, 2020). Aligned image stacks were filtered to remove noise and highlight contours using a Mean filter in Fiji (0.5 pixel radius). Segmentation of organelles (plastids, mitochondria, nucleus) and other cellular compartments of the different cell types was carried out with 3D Slicer software (Oyama *et al.*, 2014) (www.slicer.org), using a manually-curated, semi-automatic pixel clustering mode (5 to 10 slices are segmented simultaneously in z) as in (Decelle *et al.*, 2021; Uwizeye *et al.*, 2021). We assigned colors to segmented regions using paint tools and adjusted the threshold range for image intensity values. Morphometric analyses were performed with the 3D slicer module “segmentStatistics” on the different segments (segmented organelles) and converted to µm3 or µm2 taking into account the voxel size of 8 nm. In total, 5 cells were analyzed for 2n-calcified and 1n cells, 4 cells in decoupled cells that were calcified and 6 decoupled cells that were flagellated and non-calcified.

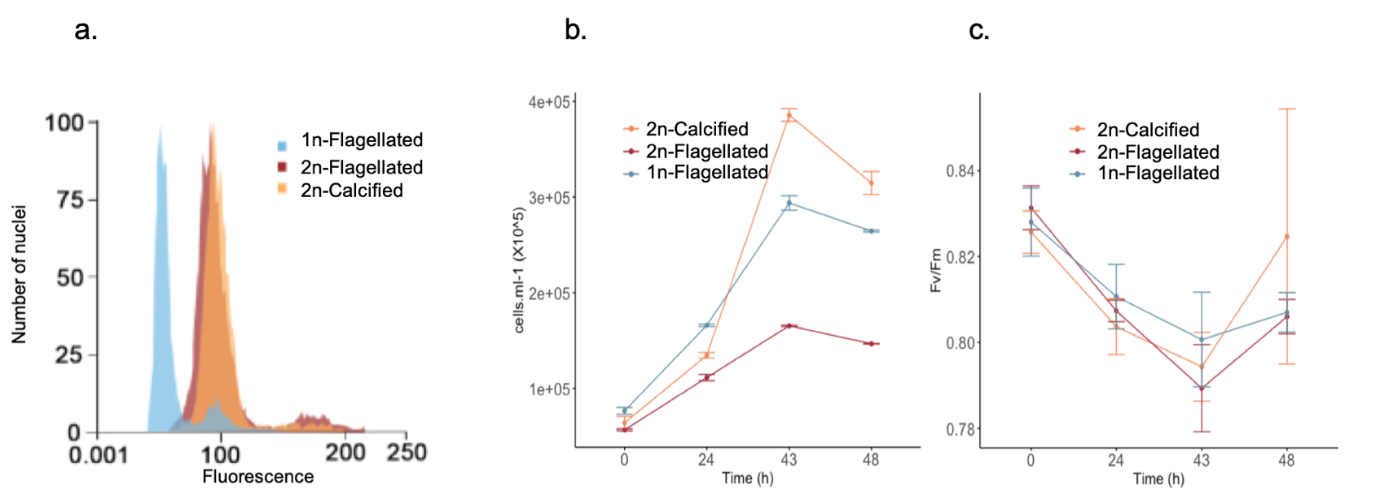

**Supporting Figure Fig. S1** Characterisation of cultures for transcriptome analysis. (a) DNA content histograms of extracted nuclei of 2n-calcified (RCC1216),1n-flagellated (LC5-11A) and 2n-flagellated (decoupled) (LC5-12A) cells at 43h (see b). Extracted nuclei were stained with Sybr Green I and analysed by flow cytometry. The main peak represents the G1 phase of the cell cycle. The high fluorescence, but minor peak the G2 phase of the cells cycle. (b) Cell concentrations in cultures at the different times of sampling. Cell counts were undertaken by flow cytometry. Samples for transcriptome analyses were collected at 43h. (c) Maximum photochemical quantum yields of PSII (Fv/Fm) of the cultures over the different times of sampling. The Fv/Fm measurements were undertaken with a fluorescence induction and relaxation system. 1n and 2n: haploid and diploid respectively.

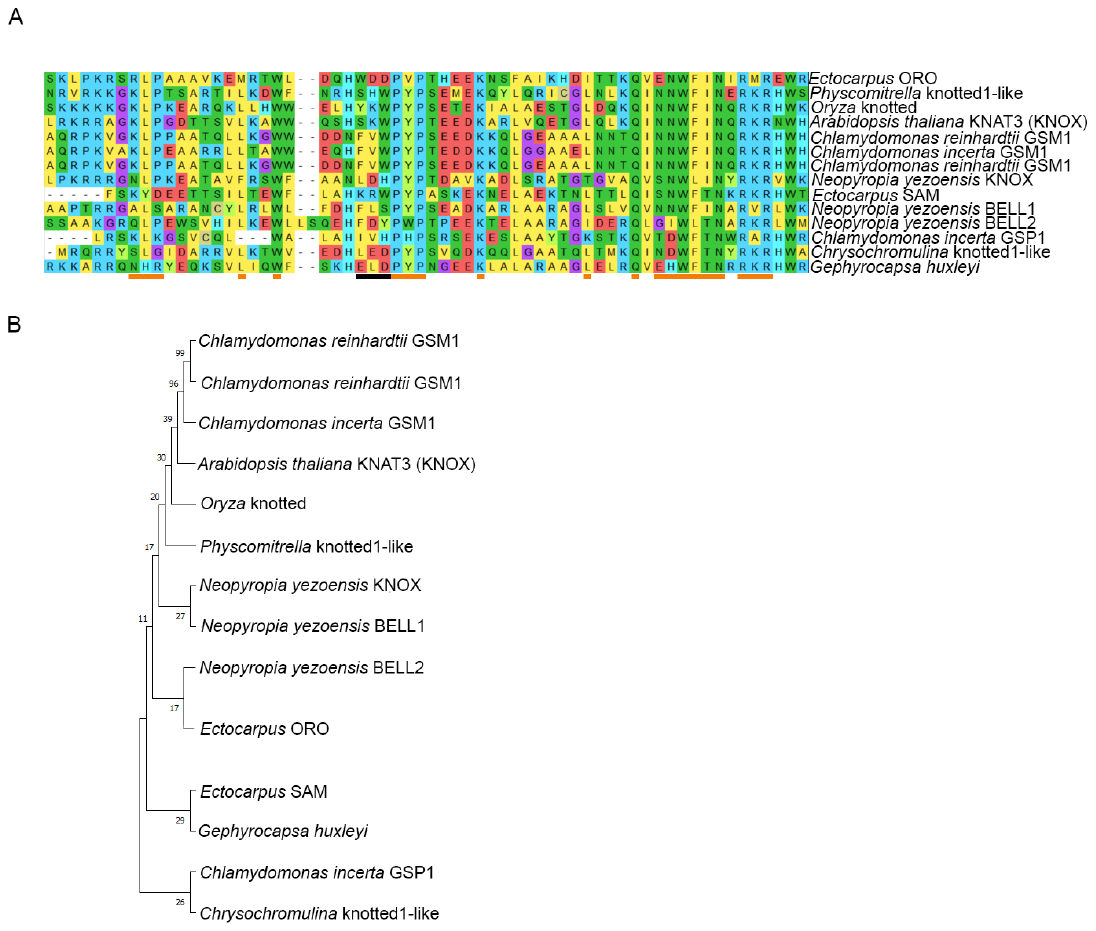

**Supporting figure Fig. S2** TALE (Three Amino acid Length Extension) Homeoproteins of plants and algae that regulate life cycle transitions. (a) Alignment of the conserved homeobox domain. The TALE is underlined with a black line. Zones of higher homology are underlined with oranges bars following Lee et al. 2008. The sequence identified of *Gephyrocapsa huxleyi* in our transcriptome dataset is c0_g13125. Accession numbers of other sequences are: *Arabidopsis thaliana*, AtKNAT3 X92392; *Neopyropia yezoensis*, MK629536, MN070241 and MN070242; *Physocmitrella*, BAF96739; *Oryza* XM015790631; *Chlamydomonas reinhardti* GSM1, ABS71849; ABJ15867; *Chlamydomonas incerta* GSM1, ABJ15868; *Chlamydomonas incerta* GSP1, AAW82031; *Ectocarpus*, KU746822 and KU746823; *Chrysochromulina* CCMP291, KOO53578.The alignment was generated in MEGA 8.0 software (https://www.megasoftware.net) using the MUSCLE algorithm. (b) Maximum Likelihood (ML)-based phylogenetic analysis of the homeobox domain. The phylogeny was constructed in in MEGA 8.0 software using LG+G substitution model and 100 bootstrap replication.
